# Neural circuit robustness to acute, global physiological perturbations

**DOI:** 10.1101/480830

**Authors:** Jacob Ratliff, Eve Marder, Timothy O’Leary

## Abstract

Neural function depends on underlying physiological processes that are highly sensitive to physical variables such as temperature. However, some robustness to perturbations in these variables manifests at the circuit level, suggesting that circuit properties are organized to tolerate consistent changes in underlying parameters. We show that a crustacean pacemaker circuit is robust to two global perturbations - temperature and pH - that differentially alter circuit properties. Consistent with high variability in underlying circuit parameters, we find that the critical temperatures and pH values where circuit activity breaks down vary widely across animals. Despite variability in critical points the network state transitions at these critical points are consistent, implying that qualitative circuit dynamics are preserved across animals, in spite of high quantitative parameter variability. Surprisingly, robustness perturbations in pH only moderately affect temperature robustness. Thus, robustness to a global perturbation does not necessarily imply sensitivity to other global perturbations.

## Introduction

All nervous systems experience fluctuations in their environment that have the potential to disrupt circuit activity. In many species, fluctuations in core physical variables such as temperature and pH are actively buffered by compensatory physiological responses and behavioral preferences (Haddad & Marder 2018, Marder et al 2015, Obara et al 2008, Pequeux 1995, Robertson & Money 2012). In addition to active mechanisms that maintain homeostasis, neural circuits exhibit intrinsic robustness to perturbations that are not compensated by other means, providing an additional line of defense against circuit failure (Roemschied et al 2014, Soofi et al 2014, Tang et al 2010, Tang et al 2012).

Recent work in crustacean nervous systems shows a core neural circuit in the Stomatogastric Ganglion (STG) of the crab, *Cancer borealis*, can maintain normal activity patterns despite very large changes in temperature spanning tens of degrees Celsius (Rinberg et al 2013, Soofi et al 2014, Tang et al 2010, Tang et al 2012). This robustness makes sense ecologically, because crustaceans such as crabs and lobsters are poikilotherms - they do not regulate their body temperature precisely - and experience natural variations in temperature in their habitat. However, all biochemical reactions are temperature-dependent, so every physiological property that underpins circuit function will be altered by a temperature change. For this reason, we refer to a temperature perturbation as a global perturbation.

There are several surprising aspects of the STG’s robustness to acute changes in temperature. The underlying physiological properties of the neurons show large and heterogeneous temperature sensitivities that differ several-fold between different currents and gating variables (Tang et al 2010). Without constraints on channel expression relationships, such strong and heterogeneous temperature dependence would detune physiological properties and cause circuit failure for modest temperature changes (Caplan et al 2014, O’Leary & Marder 2016, Robertson & Money 2012). However, it is well known that there is large (several-fold) variability in the expression of the different ionic conductances within the identified neurons of the STG (Schulz et al 2006, Schulz et al 2007). Therefore, any mechanism that tunes conductance expression to avoid temperature induced instability must also allow large variation in the space of solutions it finds. Recent work has shown how correlations in conductance expression can reconcile variability with temperature robustness, provided the correlations are constrained to offset the sensitivity of circuit behavior to channel properties (O’Leary & Marder 2016).

Together these observations suggest that robustness to temperature imposes a constraint on the physiological properties of the circuit. An immediate question is whether such a constraint might be satisfied only at the cost of making the circuit vulnerable to other kinds of global perturbations. The question we address here is how temperature robustness interacts with, or limits robustness to other global perturbations that alter circuit properties distinctly from temperature.

We investigated combined robustness of the pyloric pacemaker circuit in the STG to both temperature and pH perturbations. We subjected the same recorded neurons to simultaneous temperature and pH variations during normal ongoing circuit activity. pH has similar widespread effects on ionic currents, reversal potentials and channel kinetics as temperature (Church et al 1998, Cook et al 1984, Hille 2001, Tombaugh & Somjen 1996, Xiong & Stringer 2000), although the effects of pH on individual physiological variables in the STG are not as well characterized as those of temperature (Golowasch & Deitmer 1993). Moreover, there is some evidence that crabs and other marine organisms may experience acute changes in pH in their environment, suggesting that the circuit may be adapted to cope with this perturbation (Sartoris & Pörtner 1997, Truchot 1973, Whiteley 2011). We developed a means of quantifying internal variability in the pyloric rhythm that is predictive of the eventual collapse of the rhythm, albeit to a limited extent.

## Results

The pyloric rhythm in the stomatogastric ganglion (STG) is driven by a subset of identified neurons that comprise a so-called pacemaker kernel consisting of the two (PD, pyloric dilator) and single AB (anterior burster) neurons. The pacemaker kernel rhythmically inhibits and the LP (lateral pyloric) and PY (pyloric) neurons, as depicted in Figure 1A. The pacemaker kernel is required for a stable oscillation in the full circuit. The AB neuron is an intrinsically bursting cell that is strongly electrically coupled to the two PD neurons. The LP neuron feeds back onto and inhibits the PD neurons using glutamatergic transmission and can be seen in the intracellular waveform of the PD neuron as inhibitory post synaptic potentials (IPSPs) (Figure 1A, bottom). We focused on the pacemaker kernel, which is able to maintain a stable oscillation when isolated pharmacologically from the rest of the circuit. By adding picrotoxin (PTX), we blocked the glutamatergic transmission in the STG thereby removing the feedback connections on the pacemaking kernel (Figure 1B) (Marder & Eisen 1984). After the addition of PTX, glutamatergic IPSPs are no longer present and the membrane potential depolarizes slightly, but the activity recorded either of the PD neurons shows a stable oscillation (Figure 1B, bottom panel).

**Figure 1.**
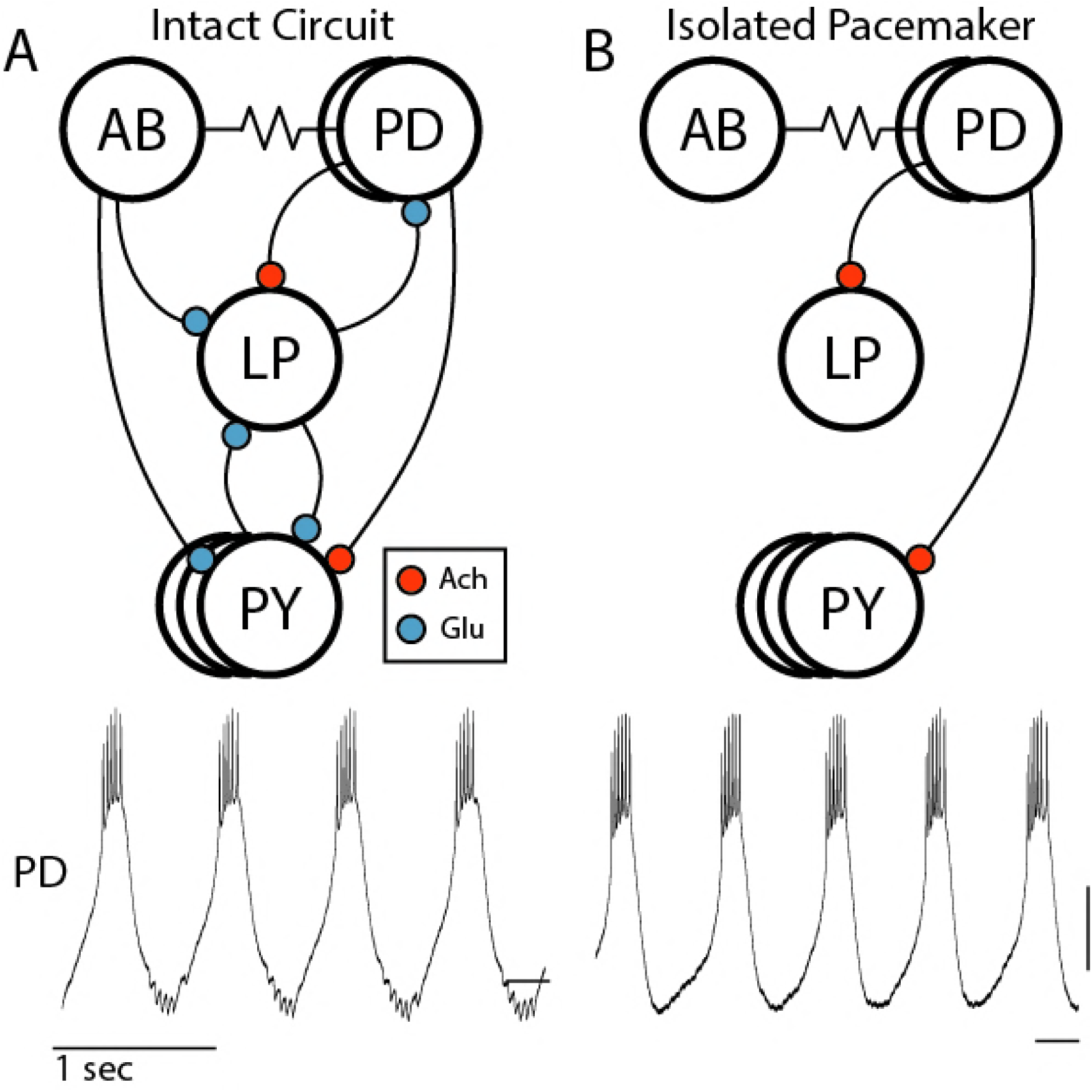
The pyloric and isolated pacemaker circuits. (A) Above: Circuit diagram of the pyloric network of the stomatogastric ganglion. Chemical synapses are represented by curved lines with colored balls where red indicates a cholinergic synapse and blue is a glutamatergic synapse. Electrical synapses are represented by resistor symbols. Below: An intracellular recording of the PD neuron from the intact pyloric circuit. (B) Above: Circuit diagram of the isolated pacemaker after the addition of PTX. Below: An intracellular recording of the PD neuron from the isolated pacemaker circuit. (A, B) Scale: 10mV with dash at −50mV

### Activity of isolated pacemaker near critical temperatures

We examined the activity of the pacemaker kernel in response to acute changes in temperature. Previous work established that the pacemaker oscillation fails at a critical temperature (Rinberg et al 2013). We roughly determined the temperature at which the intact pyloric rhythm became disorganized or silent using extracellular recordings. We denote this the *critical temperature* or *transition point*. Consistent with previous work, many preparations are robust for large changes in temperature that would prohibit a stable intracellular recording. We therefore selected preparations (13 in this set of experiments) that showed a reversible temperature-induced transition in the range 11-30°C. We then set the temperature of the bath solution to 5°C below the transition point, applied picrotoxin, and obtained intracellular recordings from a PD neuron in the isolated pacemaker. The temperature was then slowly increased at a rate of approximately 5°C per hour while holding the intracellular recording allowing us to monitor changes in the activity patterns of the isolated pacemaker with small changes in temperature (Figure 2).

**Figure 2.**
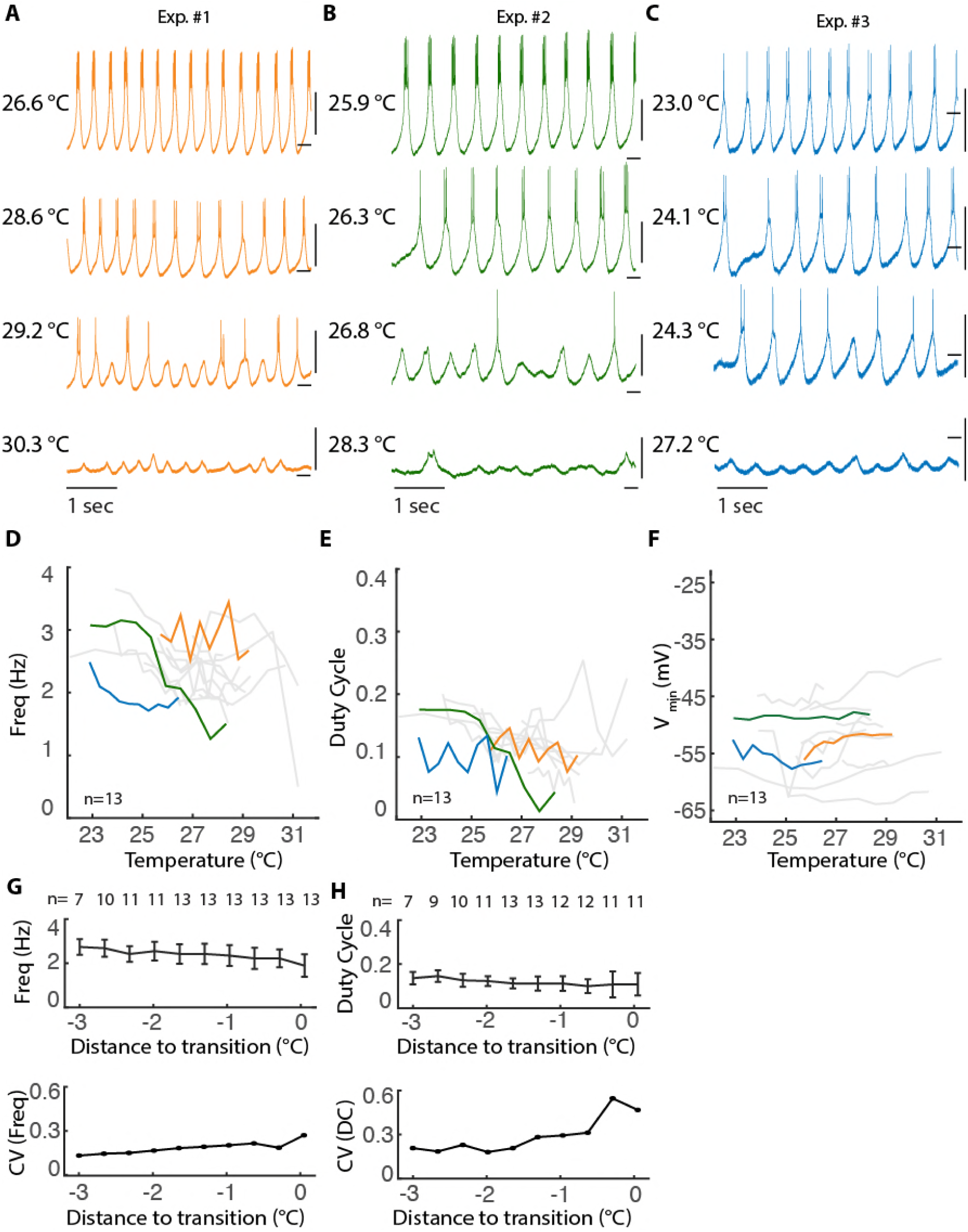
Activity of isolated pacemaker near critical temperatures. (A-C) Intracellular recordings of the PD neuron in the presence of PTX across a range of temperatures. Scale: 10mV with dash at −50mV (D-F) Burst frequency (D), duty cycle (E), and minimum voltage during oscillation (F) of PD neuron from 13 preparations plotted as a function of temperature. Duty cycle computed from intracellular traces as burst duration normalized to period of oscillation. Duty cycle becomes undefined for single spike bursts. Colored lines correspond to example experiments with same color in (A-C). (G-H) Above: average burst frequency (G) and duty cycle (H) across preparations plotted as a function of distance, in degrees Celsius, to transition to silence. Error bars represent standard deviations. Not all cells were recorded for 3 degrees before transition (see methods). Below: coefficients of variation for burst frequency and duty cycle, respectively, calculated from above plots.

Recordings from three example experiments are shown in Figures 2A-C. When far from critical temperatures, the isolated pacemaker had relatively constant burst frequency with clear membrane potential plateaus and bursts of action potentials. As temperature was increased, bursting became less regular, leaving plateaus with few or no spikes and variable interburst intervals. Qualitative changes in bursting of the pacemaker, i.e. transition points, were observed across small changes in temperature and between preparations these qualitative changes occur at different temperatures. In Figure 2A for example, there is a change in activity patterns of the preparation between 29.2°C to 30°C with bursting activity ceasing at the higher temperature. In this preparation, there is with little qualitative change between 26.6°C and 28.6°C, which can be contrasted with the changes in the preparation shown in Figure 2B where there is a dramatic change in activity pattern of the pacemaker from 26.3°C to 26.8°C. In each preparation as temperature was increased further, activity patterns transitioned to silence, with no spikes fired and only small fluctuations in the membrane potential (Figure 2A-C, bottom traces).

To quantitatively examine the changes in activity of the isolated pacemaker near critical temperatures, we measured the burst frequency, duty cycle, and minimum membrane potential plotted against absolute temperature (Figure 2D-F) and relative to the transition point to silence (Figure 2G, H). Duty cycle is defined as the duration of time spiking normalized to the burst period and has previously been shown to be conserved over temperature ranges that permit a stable oscillation (Rinberg et al 2013, Tang et al 2010).

It has previously been shown that the burst frequency of the isolated pacemaker increases with temperature between 10°C and 25°C (Rinberg et al 2013, Tang et al 2010). When examining burst frequency near critical temperatures, we found that this relationship was not present (Figure 2D) as there was no consistent increase in frequency with increasing temperature across preparations. In addition, when data were aligned to transition to silence, mean frequency across preparations decreases as preparations approach the transition, with increasing between-preparation variability (Figure 2G). Furthermore, duty cycle becomes more variable between preparations near critical temperatures (Figure 2E), and when aligned to the transitions to silence, we saw that there is greatly increased variability between preparations near critical transitions.

### Activity of isolated pacemaker near critical pH

To contrast temperature-induced changes with those induced by a second global perturbation, we examined the effects of acidic pH on the isolated pacemaker. Recent work has shown that the pyloric rhythm continues in the presence of extreme pH in an approximate range from pH 6.1 to pH 8.8 and that below approximately pH 6, the pyloric rhythm becomes silent (Haley et al 2018). We sought to examine what occurs near these critical pH levels. To do this, we obtained intracellular recordings of the PD neuron in physiological saline at 11°C (∼pH 7.8). pH was then slowly lowered by continuously adding pH 5 saline to the volume of saline feeding the bath with the rate of mixing adjusted to create a change to pH 6 over the course of one hour.

Example traces from three experiments are shown in Figures 3A-C. All preparations were bursting at pH 7 (Figure 3A-C, top traces), but as pH was deceased, the regularity of this bursting changed with preparations depolarizing and the amplitude of the slow wave decreasing (Figure 3A-C). At lower pH, bursting became intermittent with periods of tonic spiking and eventually the isolated pacemaker transitioned to tonic spiking activity (Figure 3A-C, middle). After this transition, further decreases in pH caused additional depolarization, smaller amplitude spikes, and finally the tonic spiking pattern transitioned to silence (Figure 3A-C). We therefore defined two critical pH values for each of these qualitative transitions in activity, one marking the transition from bursting to tonic spiking and one marking tonic spiking to silence.

**Figure 3.**
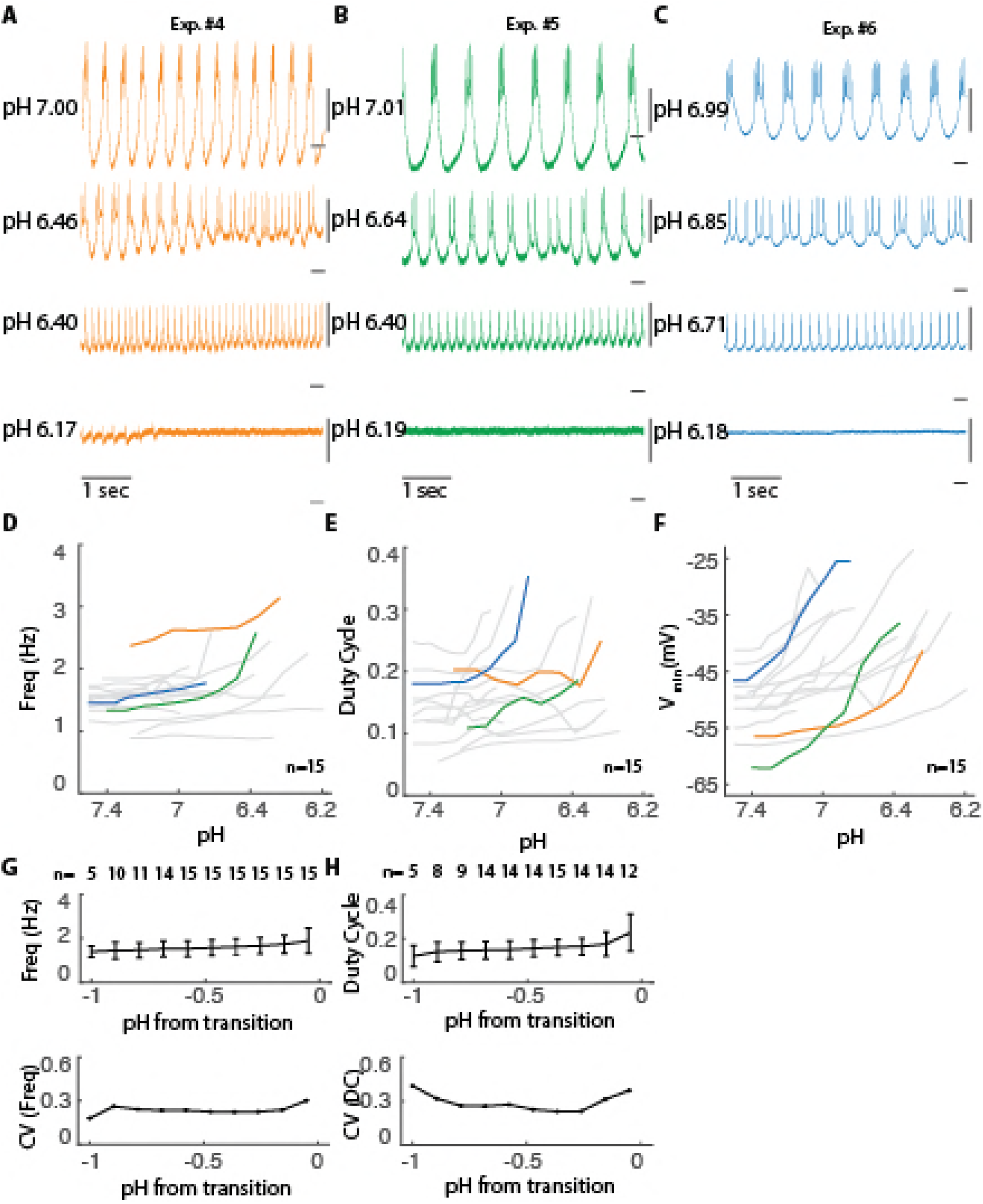
Activity of isolated pacemaker in acidic pH. (A-C) Intracellular recordings of the PD neuron in the presence of PTX across range of acidic pH. Scale: 10mV with dash at −50mV (D-F) Burst frequency (D), duty cycle (E), and minimum voltage during oscillation (F) of PD neuron plotted as a function of pH from 15 preparations. Duty cycle computed as in figure 2. Colored traces correspond to example experiments of same color in (A-C). Frequency and duty cycle are only plotted when cell is bursting. (G) Above: average burst frequency (G) and duty cycle (H) across preparations plotted as a function of distance to transition to tonic spiking. Error bars represent standard deviations. Not all cells were recorded for a range of 1 pH. Below: coefficients of variation for burst frequency and duty cycle, respectively, calculated from above plot.

To compare the effects of pH and temperature near critical points (Figure 2D-H), we examined the relationship of pH with burst frequency, duty cycle, and minimum voltage during oscillations (burst frequency and duty cycle are only defined during bursting). In the range examined here, there is little effect of pH on burst frequency with a modest trend toward increased burst frequency (Figure 3D, G). Duty cycle near critical pH is variable, with many, but not all, preparations having increased duty cycle prior to transition to tonic spiking (Figure 3E, H). In contrast preparations near critical temperatures (Figure 2F), changes in pH cause substantial depolarization (Figure 3F). The qualitative and quantitative differences between pH- and temperature-induced changes in membrane potential activity are consistent with these perturbations having distinct, global effects on underlying membrane currents, as shown in previous work (Church et al 1998, Doering & McRory 2007, Golowasch & Deitmer 1993, Tombaugh & Somjen 1996).

### Predicting transitions in isolated pacemaker

We have shown that near critical temperatures and pH the activity patterns of the isolated pacemaker change abruptly with critical points varying between preparations. These transitions occur at different temperatures and pH in different preparations. There is a body of theory (Chisholm & Filotas 2009, Kandel et al 1988, Scheffer 2010, Scheffer et al 2012, Veraart et al 2012) that proposes a set of generalized markers to predict critical transitions in dynamical systems, including complex biological systems. These markers include increased variability, increased recovery time from perturbation, and flickering between states. We therefore analyzed membrane potential variability near transitions to assess the power to predict the precise transition points in the activity patterns of the isolated pacemaker.

Increased variability is depicted in Figure 4A and B using an example system consisting of a ball in a trough, subject to noisy perturbations. This system is stably attracted to state 0, the lowest energy state, while noise moves the balls randomly away from this stable point. As the system moves closer to reorganizing, thereby gaining a new stable state, the basin of attraction shallows (Figure 4B). The same amount of noise now generates greater variation in the movement of the ball. This simple example illustrates why increased variability is expected near a transition point in a dynamical system: ongoing, internal noise perturbations cause variability in the system’s dynamics. As the system approaches a transition, its sensitivity generically increases, and the impact of the internal noise becomes more visible.

**Figure 4.**
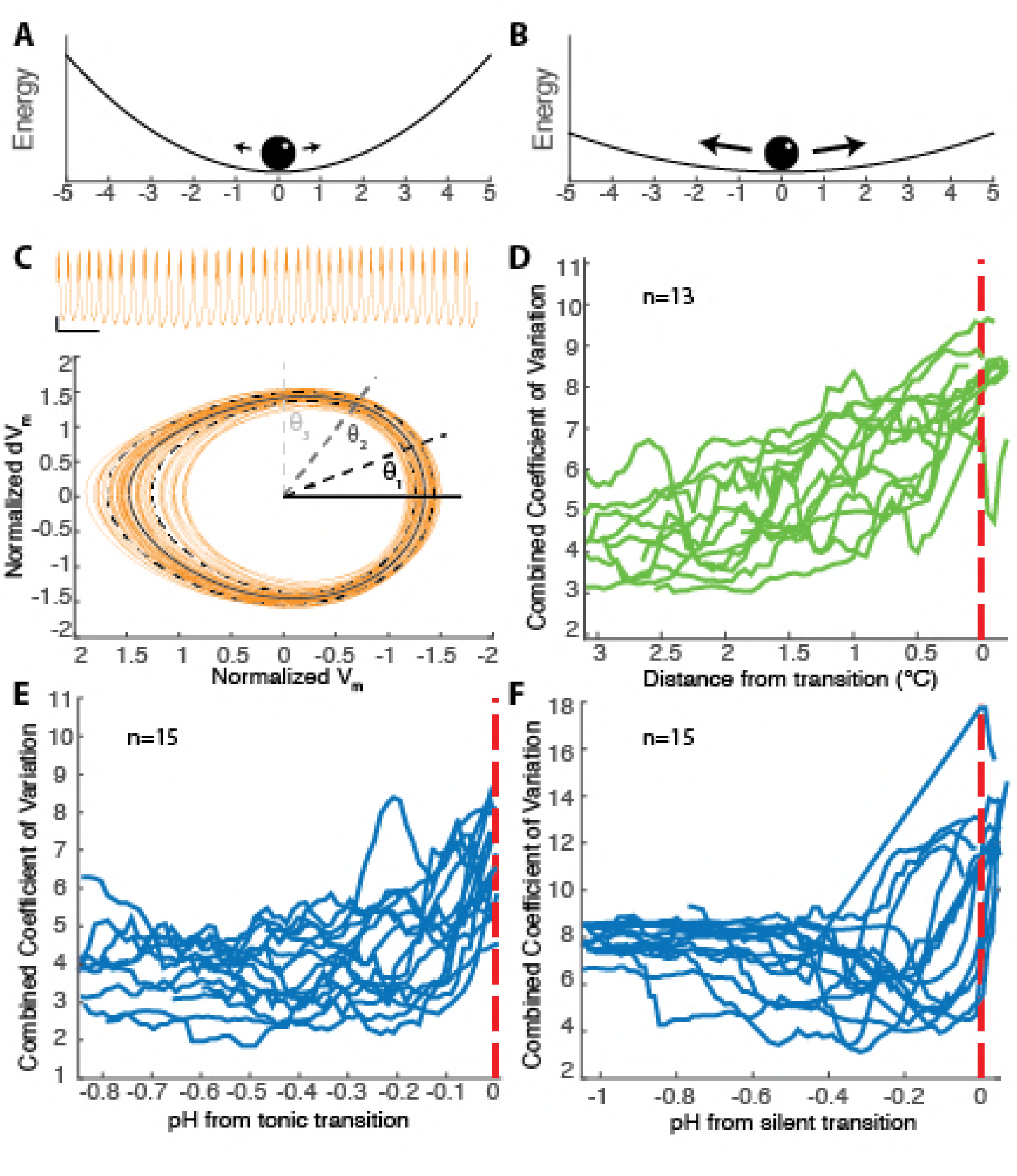
Variance increases at the population level prior to transitions in activity pattern. (A, B) Cartoon schematic depicting noisy ball attracted to bottom of trough. The same amount of noise moves the ball more in (B) compared to (A). (C) Above: voltage trace from PD neuron in isolated pacemaker plotted in orange. Scale: 1 second, 5mV at −50mV. Below: Phase portrait generated from low passed voltage trace plotted in orange as normalized membrane voltage (V_m_) versus normalized instantaneous change in voltage (dV_m_, see methods). The solid black line represents the mean of the oscillations and the dashed black lines are two standard deviations plus and minus the mean. These values, means and standard deviations, are calculated for 200 points in phase schematized by the solid and dashed black and grey lines. (D) Each green line represents the moving average of combined coefficients of variation (see methods) plotted as a function of temperature from transition to silence (red line). (E-F) Each blue line represents the moving average of combined coefficient of variation as a function of pH. Experiments are aligned to transition. (E) Red line represents transition to tonic spiking. (F) Red line represents transition to silence.

W examined within-preparation variance as a predictor of transitions by examining the membrane potential traces in their phase plane, as shown in Figure 4C. This allowed us to define the ‘mean oscillation,’ by computing the mean trajectory across multiple oscillations, and a coefficient of variation (CV, standard deviation normalized to the mean). This provides a measure of the internal variability of the oscillation from its average trajectory. We then combined these CV values (see methods) to compute an overall measure of variability (Combined Coefficient of Variation, CCV) and plotted this as a function of distance to a transition in both temperature and pH-induced transitions. These CCV values are shown in Figure 4 (D-F), aligned to respective transition points (dashed red line).

We analyzed variability in all preparations near temperature and pH-induced transitions. Consistent with theoretical predictions, there was a general trend for the CCV to increase near a transition. Importantly this increase occurs irrespective of the type of transition or the perturbation (temperature or pH) that led to it. However, this measure offers a poor prediction of proximity to a transition within any given preparation. For example, with the temperature perturbation, a CCV value of 6 could mean the preparation is at the transition point or more than 3 degrees away. In the case of transition to silence due to pH perturbation, the variance in many of the preparations decreases near the transition to silence. Thus, while variability at the population level shows a robust increase near transition points, there is large inter-preparation variability in this relationship that would preclude its use as a predictive tool for the onset of a transition in any given preparation.

### Combined effects of temperature and pH

Lastly, we sought to understand the relationship between pH perturbations and temperature perturbations. We started by performing pH perturbations at 25°C, obtaining intracellular recordings of the PD neuron in the isolated pacemaker, a temperature at which all preparations are bursting (Figure 5A). We then subjected these same preparations to decreasing pH ramps. The transition points are highly variable across preparations; as a consequence, pH ramps performed at 11°C and 25°C show transitions in overlapping ranges (Figure 5A, C).

**Figure 5.**
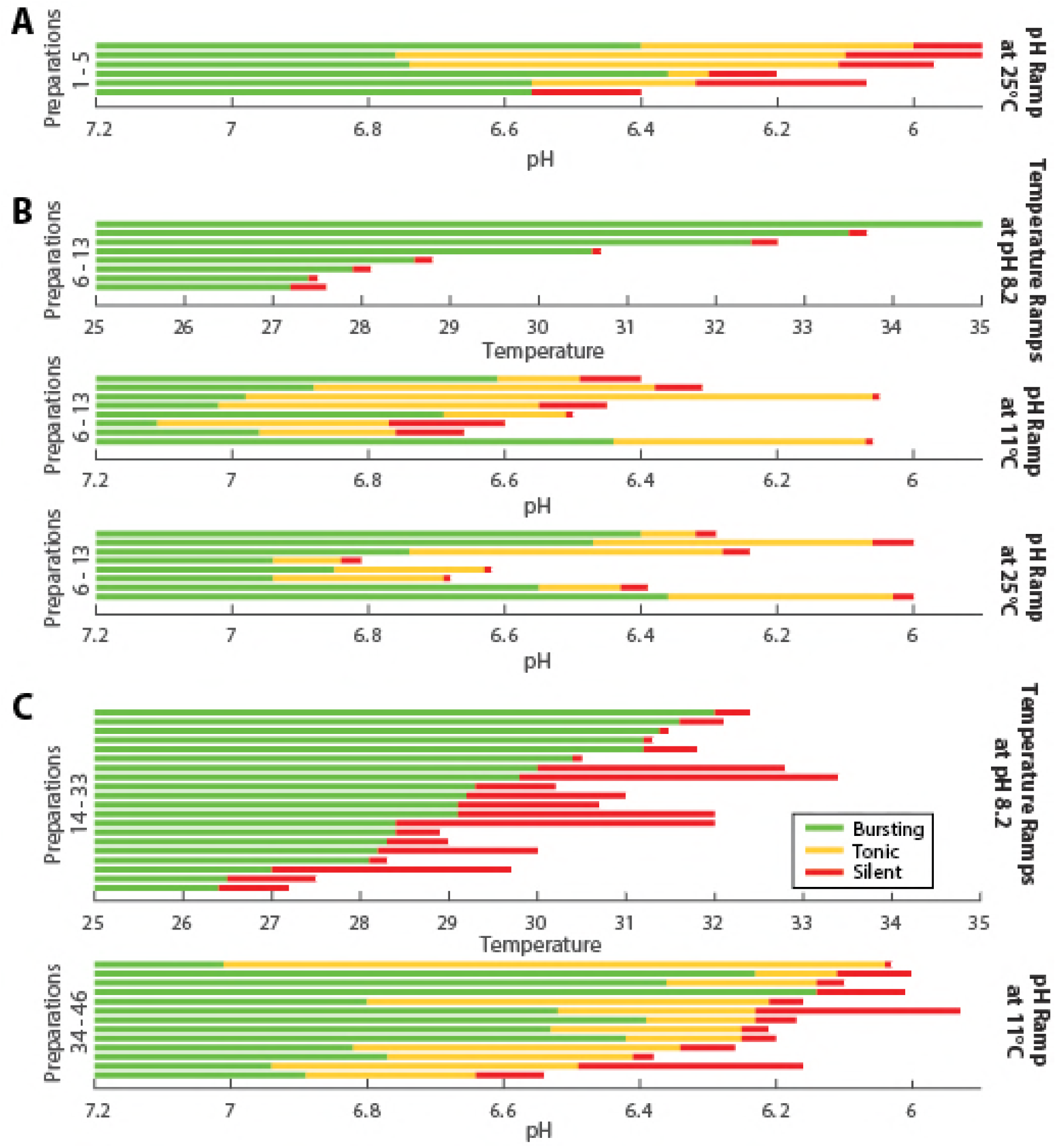
Stereotyped activity patterns across temperature and pH. (A) Each horizontal line represents one preparation exposed to a range of pH at 25°C (n=5). The qualitative activity pattern, or state, is indicated with color, green corresponding to bursting, yellow to tonic spiking, and red to silence. (B) The same preparation was exposed to each condition and plotted in the same order across conditions (n=8). Meaning the first horizontal line the temperature condition corresponds to the first horizontal line in the pH conditions. (C) The top set of preparations were exposed to increasing temperature (n=20, 13 from figure 2 and 7 additional without intracellular recordings) and the bottom set of preparations were exposed to decreasing pH (n=15) (A-C) Preparations are ordered based on transition to silence.

To control for inter-preparation variability when testing the interaction of pH and temperature, we exposed preparations to multiple perturbations: decreasing pH at 11°C, decreasing pH at 25°C, and increasing temperature in a set of seven preparations (Figure 5B). This allowed us to test two hypotheses: that the combination of temperature and pH will make preparations more sensitive (transition at less extreme values) or that preparations may be ‘tuned’ for robustness to one perturbation over another (preparations more sensitive to pH will be less sensitive to temperature and vice versa). Surprisingly, neither of these possibilities holds true in the data. At more extreme temperatures, preparations transitioned to tonic spiking at more extreme pH at 25°C compared to their transition points at 11°C. Together, these results show that there is a modest interaction between temperature and pH perturbations which, surprisingly confers slightly higher pH robustness at more extreme temperatures.

### Stereotyped transitions during temperature and pH perturbations

We have shown that the pacemaker oscillation undergoes different types of transitions in activity patterns when exposed to temperature and pH respectively. In Figure 5C, we plotted the activity patterns as a function of temperature of pH, respectively for the set of experiments from Figures 2 and 3. In each of the temperature experiments combined with those from Figure 5B, all 26 preparations transitioned from bursting to silence without tonic spiking. In contrast, 25 of 26 pH experiments transitioned from bursting to tonic spiking to silence while the remaining one transitioned from bursting to silence.

## Discussion

As a central pattern generator, the STG is well known for its reliable and stereotyped behavior. Numerous studies have shown that the STG is robust to physiological insults and pharmacological manipulation. Such robustness is essential, given the importance of the circuit’s function for the animal’s survival. Paradoxically, the reliable and stereotyped output of the STG belies the complexity and variability of the physiological mechanisms that ultimately govern circuit behavior. Ion channel densities are highly variable, with each preparation having its own unique configuration that somehow gives rise to a reliable and robust rhythm (Goaillard et al 2009, Grashow et al 2010, Haddad & Marder 2018, O’Leary et al 2013, O’Leary et al 2014, Robertson & Money 2012, Schulz et al 2006, Schulz et al 2007, Taylor et al 2009, Temporal et al 2014).

We have shown that a key subcircuit in the STG - the pyloric pacemaker kernel - is simultaneously robust to two different global perturbations. Temperature and pH were acutely covaried over a large range without disrupting the pacemaker rhythm. This suggests that in spite of animal to animal variability, the circuit has found parameters that allow detuning of ionic currents and synaptic properties to occur, but nonetheless ensure a stable rhythmic output. Computational studies show that this is a far from trivial result (Caplan et al 2014, Roemschied et al 2014). The region of functional parameter space occupied by a circuit is relatively small. Moreover, for the circuit to remain robust to temperature and pH perturbations that cause parameters to change significantly it is clear that the biological mechanisms which tune circuit properties do so in a way that ensure specific functional organization between physiological parameters amid a large degree of variability (O’Leary & Marder 2016).

We found that the pyloric pacemaker circuit is remarkably robust to acute pH variations. This robustness is somewhat dependent on temperature, indicating that both kinds of robustness impose constraints on channel expression. However, the interaction between robustness to temperature and pH robustness was surprisingly small. This implies that the circuit occupies a region of physiological parameter space that allows temperature and pH robustness be satisfied without a severe tradeoff, as well as allowing large internal variability in ionic current expression.

Consistent with underlying parameter variability, we find that pH and temperature cause the pacemaker oscillation to fail at critical values of temperature and pH that vary significantly between animals. Importantly, the modes of failure correspond to reversible transitions to distinct activity regimes, from bursting to tonic spiking and then silence in the case of a pH ramp, and from bursting to silence in the case of a temperature ramp. In agreement with general theory of critical transitions in dynamical systems we detect an increase in intrinsic variability of the oscillator close to the critical point at which the oscillation fails (Chisholm & Filotas 2009, Scheffer 2010, Scheffer et al 2012, Veraart et al 2012). The consistency of these qualitative transitions between preparations is strong evidence that the pyloric circuit operates with a consistent type of oscillatory dynamics. Together, these findings show that while large variability is indeed present in the physiological properties of the STG, the mechanisms that organize physiological parameters place the circuit in a highly robust regime with consistent qualitative behavior. This suggests that the circuit doesn’t merely achieve a robust oscillation, it achieves the same qualitative type of oscillation in spite of large variability in underlying physiological variables.

In spite of the surprising robustness we have characterized in this circuit, there is clear evidence of underlying parameter variability. Although most preparations undergo the same transitions between different activity patterns as pH and temperature are varied, the precise values at which these transitions occur is variable. On the other hand, the transitions between different activity patterns were remarkably reliable: temperature elevation consistently resulted in a transition from bursting to silence, while in most preparations a decrease in pH resulted in a sequence of transitions from bursting to tonic spiking, then from tonic spiking to silence. Together, these findings illustrate that collective circuit properties can be highly consistent, even if quantitative, low-level parameters are not. A plausible explanation for how such consistency arises is that cellular components such as ion channels are regulated in a collective, modular fashion, with multiple channel types co-regulated by the same molecular pathway (O’Leary & Marder 2016, O’Leary et al 2014, Temporal et al 2014).

We can view the pacemaker kernel preparation as a vastly simplified biological model of a nervous system that is subject to particular failure modes. Other more complex nervous systems such as the brains of vertebrate species exhibit many more components and kinds of behavior, but they also show stereotyped failure modes such as seizures. Our findings illustrate just how difficult it is to predict the onset of failure, even with what might be considered ideal biological replicates of the same system. We found that at the population level, increases in rhythm variance were indicative of the proximity to a transition out of the rhythm. This is consistent with recent theory (Chisholm & Filotas 2009, Scheffer 2010, Scheffer et al 2012) and experimental attempts to predict catastrophic events in complex natural systems (Veraart et al 2012). However, in our data the trend in variance is far from predictive at the individual level.

Our main motivation for studying combined global perturbations to a neural circuit was to assess whether robustness to one kind of perturbation implied sensitivity to other kinds of perturbations. For pH and temperature perturbations in the STG, we find a surprisingly modest interaction in the robustness of the pacemaker rhythm. This suggests that the circuit may have evolved to exhibit tolerance to both (and likely other) external insults. This combined tolerance places additional constraints on the expression and regulation of the underlying membrane currents and synaptic connections (O’Leary & Marder 2016), and may even favor specific kinds of circuit architectures over others.

## Methods

### Animals

*Cancer borealis* were purchased from Commercial Lobster (Boston, MA) and maintained at 11°C in tanks containing artificial seawater. Animals used in this study were obtained between July 2016 and November 2017.

### Solutions

*C. borealis* physiological saline was composed of 440mM NaCl, 26mM MgCl_2_, 13mM CaCl_2_, 11mM KCl, 12mM Trizma Base, and 5mM maleic acid, pH 7.4-7.5 (measured at room temperature). For more acidic saline, pH was adjusted with additional maleic acid. Picrotoxin (PTX) was purchased from Sigma (St Louis, MO) and used at 10^-5^ M in physiological saline. The microelectrode solution was 10mM MgCl_2_, 400mM KGluconate, 10mM Hepes, 15mM NaSO_4_, 20mM NaCl, pH 7.45. (Hooper et al 2015)

### Electrophysiology

The stomatogastric nervous system was dissected from the animal and pinned taught in a Sylgard (Dow Corning, Midland, MI) coated plastic Petri dish containing chilled physiological saline. All preparations used had intact inferior and superior esophageal nerves and included commissural and esophageal ganglia. For the duration of experiments, the dish was superfused with saline. Temperature was controlled using a Peltier device (Warner Instruments) and monitored using a thermistor probe placed in the dish.

Vaseline wells were placed around the lateral ventricular nerve (*lvn*) and the pyloric dilator nerve (*pdn*) and extracellular recordings were obtained using stainless steel pin electrodes placed in the wells and amplified using a differential amplifier (A-M Systems, Sequim, WA). In addition, intracellular recordings were obtained from the pyloric dilator (PD) somata using 15-25 MΩ glass microelectrodes pulled with a Flaming/Brown micropipette puller (Sutter Instrument Company, Novato, CA). The cell type was identified by comparing spiking activity to extracellular recordings on the *pdn* and by examining the intracellular waveform.

#### Temperature Manipulations

Intracellular recordings were begun at either 25°C or 7 degrees below a ‘crash’ temperature determined with extracellular recordings. Preparations were then exposed to continuously increasing temperatures, referred to as temperature ramps. A waveform generator (Rigol, Beijing, China) was used to create a steadily increasing voltage to control the output of the Peltier device. Temperature was increased until preparations changed from bursting to silence without continuous bursting/spiking activity at which point the temperature ramp was stopped in a majority of experiments.

As previously reported, somata swelled with increasing temperature (Rinberg et al 2013, Tang et al 2012). With increasing temperature, small adjustments to the location of the intracellular electrode were made to maintain the recording.

All preparations analyzed, with the exception of the experiments shown in Figure 5B, were selected based on the presence of the transition to silence. Preparations that continued to burst past 34°C were excluded.

#### pH Manipulations

Intracellular recordings of the PD neuron were begun at physiological pH. After the addition of PTX, the pH of the superfused saline was controlled by a slow, continuous mixing of pH 7 and pH 5 physiological saline during the experiments. The pH of the superfused saline was measured using a pH microelectrode purchased through Thermo Scientific (Orion 9810BN; Waltham, MA). The probe was calibrated each day using reference solutions at 11°C and/or 25°C.

### Data Analysis

Data were acquired using a Digidata 1440 data acquisition board (Axon Instruments, San Jose, California) and analyzed using MATLAB (MathWorks, Natick, MA).

#### Transition Definitions

Discrete transition points were defined in both the pH and temperature experiments with similar definitions used in both. A transition was marked when a preparation spent more than 20 seconds out of 30 second period in any activity pattern. This reliably captured switches from one activity pattern to another while filtering out small flickering events between activity patterns that occur in small ranges (<0.5°C or <0.2 pH) near transitions.

#### Phase plane analysis

The stretches of data to be analyzed were first low-pass filtered to remove spikes. The filtered voltage signal and its derivative were normalized by the standard deviation of each respective signal. The signal was mean subtracted to center the oscillation on the origin of the axes and then transformed from a Cartesian coordinate system (with the normalized voltage signal on the x-axis and the normalized voltage derivative on the y-axis) to polar coordinates. These steps generate the phase portrait shown in Figure 4A.

Next, we calculated the average trajectory of the oscillations by taking the mean and standard deviation of the radial coordinate at 200 evenly spaced angular coordinates. This gave use the envelope plotted in the phase plane in Figure 4A. From these values, we calculated the coefficient of variation, the standard deviation normalized to the mean, at each point in the phase of the oscillation. We then combined the values by taking their root-mean-square and these values were plotted in Figure 4B-D after being smoothed by taking a 0.5°C or 0.1 pH moving average.

## Acknowledgements

We would like to acknowledge Anatoly Rinberg for his contribution to this work in performing experiments that were preliminary to the study here. We would also like to thank Jessica Haley for sharing experimental results allowing for the design of these experiments. This work is funded by NIH grant R35 NS 097343-03 to E.M.

## References

Caplan JS, Williams AH, Marder E. 2014. Many parameter sets in a multicompartment model oscillator are robust to temperature perturbations. The Journal of neuroscience : the official journal of the Society for Neuroscience 34: 4963–75

Chisholm RA, Filotas E. 2009. Critical slowing down as an indicator of transitions in two-species models. Journal of theoretical biology 257: 142–49

Church J, Baxter KA, McLarnon JG. 1998. pH modulation of Ca2+ responses and a Ca2+-dependent K+ channel in cultured rat hippocampal neurones. The Journal of physiology 511: 119–32

Cook DL, Ikeuchi M, Fujimoto WY. 1984. Lowering of pHi inhibits Ca2+-activated K+ channels in pancreatic B-cells. Nature 311: 269

Doering C, McRory J. 2007. Effects of extracellular pH on neuronal calcium channel activation. Neuroscience 146: 1032–43

Goaillard J-M, Taylor AL, Schulz DJ, Marder E. 2009. Functional consequences of animal-to-animal variation in circuit parameters. Nature neuroscience 12: 1424

Golowasch J, Deitmer J. 1993. pH regulation in the stomatogastric ganglion of the crab Cancer pagurus. Journal of Comparative Physiology A 172: 573–81

Grashow R, Brookings T, Marder E. 2010. Compensation for variable intrinsic neuronal excitability by circuit-synaptic interactions. Journal of Neuroscience 30: 9145–56

Haddad SA, Marder E. 2018. Circuit Robustness to Temperature Perturbation Is Altered by Neuromodulators. Neuron 100: 609–23 e3

Haley JA, Hampton D, Marder E. 2018. Two central pattern generators from the crab, Cancer borealis, respond robustly and differentially to extreme extracellular pH. BIORXIV 2018/374405

Hille B. 2001. Ion channels of excitable membranes. Sinauer. xviii, 814 p., [8] p. of plates pp.

Hooper SL, Thuma JB, Guschlbauer C, Schmidt J, Buschges A. 2015. Cell dialysis by sharp electrodes can cause nonphysiological changes in neuron properties. J Neurophysiol 114: 1255–71

Kandel D, Domany E, Ron D, Brandt A, Loh Jr E. 1988. Simulations without critical slowing down. Physical review letters 60: 1591

Marder E, Eisen JS. 1984. Transmitter identification of pyloric neurons: electrically coupled neurons use different neurotransmitters. J. Neurophysiol. 51: 1345–61

Marder E, Haddad SA, Goeritz ML, Rosenbaum P, Kispersky T. 2015. How can motor systems retain performance over a wide temperature range? Lessons from the crustacean stomatogastric nervous system. Journal of Comparative Physiology A 201: 851–56

O’Leary T, Marder E. 2016. Temperature-Robust Neural Function from Activity-Dependent Ion Channel Regulation. Curr Biol 26: 2935–41

O’Leary T, Williams AH, Caplan JS, Marder E. 2013. Correlations in ion channel expression emerge from homeostatic tuning rules. Proceedings of the National Academy of Sciences of the United States of America 110: E2645–54

O’Leary T, Williams AH, Franci A, Marder E. 2014. Cell types, network homeostasis, and pathological compensation from a biologically plausible ion channel expression model. Neuron 82: 809–21

Obara M, Szeliga M, Albrecht J. 2008. Regulation of pH in the mammalian central nervous system under normal and pathological conditions: facts and hypotheses. Neurochemistry international 52: 905–19

Pequeux A. 1995. Osmotic regulation in crustaceans. Journal of Crustacean Biology 15: 1–60

Rinberg A, Taylor AL, Marder E. 2013. The effects of temperature on the stability of a neuronal oscillator. PLoS computational biology 9: e1002857

Robertson RM, Money TG. 2012. Temperature and neuronal circuit function: compensation, tuning and tolerance. Curr Opin Neurobiol 22: 724–34

Roemschied FA, Eberhard MJ, Schleimer JH, Ronacher B, Schreiber S. 2014. Cell-intrinsic mechanisms of temperature compensation in a grasshopper sensory receptor neuron. Elife 3: e02078

Sartoris F-J, Pörtner H-O. 1997. Temperature dependence of ionic and acid-base regulation in boreal and arctic Crangon crangon and Pandalus borealis. Journal of experimental marine biology and ecology 211: 69–83

Scheffer M. 2010. Complex systems: foreseeing tipping points. Nature 467: 411

Scheffer M, Carpenter SR, Lenton TM, Bascompte J, Brock W, et al. 2012. Anticipating critical transitions. science 338: 344–48

Schulz DJ, Goaillard JM, Marder E. 2006. Variable channel expression in identified single and electrically coupled neurons in different animals. Nat Neurosci 9: 356 – 62

Schulz DJ, Goaillard JM, Marder EE. 2007. Quantitative expression profiling of identified neurons reveals cell-specific constraints on highly variable levels of gene expression. Proceedings of the National Academy of Sciences of the United States of America 104: 13187–91

Soofi W, Goeritz ML, Kispersky TJ, Prinz AA, Marder E, Stein W. 2014. Phase maintenance in a rhythmic motor pattern during temperature changes *in vivo*. J Neurophysiol 111: 2603–13

Tang L, Goeritz M, Caplan J, Taylor A, Fisek M, Marder E. 2010. Precise Temperature Compensation of Phase in a Rhythmic Motor Pattern. PLoS Biol 8: e1000469

Tang LS, Taylor AL, Rinberg A, Marder E. 2012. Robustness of a rhythmic circuit to short- and long-term temperature changes. The Journal of neuroscience : the official journal of the Society for Neuroscience 32: 10075–85

Taylor AL, Goaillard JM, Marder E. 2009. How multiple conductances determine electrophysiological properties in a multicompartment model. The Journal of neuroscience : the official journal of the Society for Neuroscience 29: 5573–86

Temporal S, Lett KM, Schulz DJ. 2014. Activity-dependent feedback regulates correlated ion channel mRNA levels in single identified motor neurons. Curr Biol 24: 1899–904

Tombaugh GC, Somjen GG. 1996. Effects of extracellular pH on voltage-gated Na+, K+ and Ca2+ currents in isolated rat CA1 neurons. The Journal of physiology 493: 719–32

Truchot J. 1973. Temperature and acid-base regulation in the shore crab Carcinus maenas (L.). Respiration physiology 17: 11–20

Veraart AJ, Faassen EJ, Dakos V, van Nes EH, Lürling M, Scheffer M. 2012. Recovery rates reflect distance to a tipping point in a living system. Nature 481: 357

Whiteley N. 2011. Physiological and ecological responses of crustaceans to ocean acidification. Marine Ecology Progress Series 430: 257–71

Xiong Z-Q, Stringer JL. 2000. Extracellular pH responses in CA1 and the dentate gyrus during electrical stimulation, seizure discharges, and spreading depression. Journal of Neurophysiology 83: 3519–24

